# Microsecond melting and revitrification of cryo samples with a correlative light-electron microscopy approach

**DOI:** 10.1101/2022.09.14.507962

**Authors:** Gabriele Bongiovanni, Oliver F. Harder, Marcel Drabbels, Ulrich J. Lorenz

## Abstract

We have recently introduced a novel approach to time-resolved cryo-electron microscopy (cryo-EM) that affords microsecond time resolution. It involves melting a cryo sample with a laser beam to allow dynamics of the embedded particles to occur. Once the laser beam is switched off, the sample revitrifies within just a few microseconds, trapping the particles in their transient configurations, which can subsequently be imaged to obtain a snap shot of the dynamics at this point in time. While we have previously performed such experiments with a modified transmission electron microscope, we here demonstrate a simpler implementation that uses an optical microscope. We believe that this will make our technique more easily accessible and hope that it will encourage other groups to apply microsecond time-resolved cryo-EM to study the fast dynamics of a variety of proteins.

## Introduction

Structure determination of proteins has made rapid progress in the last decade, particularly thanks to the resolution revolution in cryo-EM (Kühlbrandt, 2014; Nakane *et al*., 2020; Yip *et al*., 2020), which now appears set to become the preferred method in structural biology (Hand, 2020), and the advent of machine learning approaches for protein structure prediction (Baek *et al*., 2021; Jumper *et al*., 2021). At the same time, these advances put into relief our incomplete understanding of the dynamics and the function of proteins (Henzler-Wildman and Kern, 2007). In fact, it has been argued that understanding and ultimately predicting protein function is the next frontier in structural biology (Ourmazd *et al*., 2022).

Our incomplete understanding of protein function is to a large extent a consequence of the difficulty of observing proteins as they perform their task. This requires not only near-atomic spatial resolution, but also a time resolution that is sufficient to observe the domain motions that are frequently associated with the activity of a protein and that typically occur on short timescales of microsecond to milliseconds (Boehr *et al*., 2006; Henzler-Wildman and Kern, 2007). Time-resolved cryo-EM has enabled observations of a range of processes (Frank, 2017; Fu *et al*., 2019; Carbone *et al*., 2021). Typically, dynamics are initiated by rapidly mixing two reactants and spraying them onto a specimen grid, which is then rapidly plunge frozen to trap short-lived intermediates (Berriman and Unwin, 1994; Frank, 2017; Kontziampasis *et al*., 2019; Dandey *et al*., 2020; Klebl *et al*., 2020). However, the time resolution of this method is fundamentally limited by the time required for plunge freezing, which is on the order of one millisecond (Frank, 2017), too slow to observe many relevant dynamics.

We have recently introduced a novel approach to time-resolved cryo-EM that affords microsecond time resolution and is thus fast enough observe many domain motions (Voss *et al*., 2021b, 2021a; Harder *et al*., 2022). We employ a laser beam to locally melt a cryo sample for several tens of microseconds, providing a well-defined time window during which the proteins can undergo conformational motions in liquid. A range of stimuli is conceivable that can be used to initiate specific dynamics. For example, caged compounds can be used to release ATP, ions, small peptides, or induce a pH jump (Shigeri *et al*., 2001; Ellis-Davies, 2007).

As the dynamics of the particles unfold, the heating laser is switched off, so that the sample rapidly cools and revitrifies, arresting the particles in their transient configurations.

We have demonstrated the viability of our approach and characterized the spatial and temporal resolution it affords. Proof-of-principle experiments confirm that once the sample is laser melted, particles can undergo motions in liquid and that upon revitrification, we can trap them in their transient states with microsecond time resolution (Voss *et al*., 2021b, 2021a). The success of a revitrification experiment can be assessed on the fly. In a successful experiment, the revitrified area in the center of the laser focus is surrounded by a region in which the sample has crystallized since its temperature has not exceeded the melting point. By adjusting the laser power to keep the diameter of the revitrified area constant, one can thus ensure that in each experiment, the sample undergoes the same temperature evolution (Voss *et al*., 2021a). We have also demonstrated that the melting and revitrification process leaves the proteins intact (Harder *et al*., 2022). Near-atomic resolution reconstructions can be obtained from revitrified cryo samples, suggesting that the revitrification process does not fundamentally limit the obtainable spatial resolution (manuscript in preparation; see also Lorenz, U. J., 2022). With all crucial aspects of our methods established, we have recently also begun to apply it to study the fast dynamics of a variety of systems.

By enabling atomic-resolution observations of the microsecond dynamics of proteins, our method promises to fundamentally advance our understanding of protein function. However, for our technique to achieve this goal, it is crucial to ensure that it is easily accessible, so that a large number of groups can adopt it. We currently perform our melting and revitrification experiments *in situ*, using a transmission electron microscope that we have modified for time-resolved experiments (Olshin *et al*., 2020). Setting up such an instrument, which few labs have at their disposal, presents a significant hurdle for the adoption of our technique. It is therefore desirable to develop a technically less involved variant.

Here, we present a simple implementation of our technique that uses an optical microscope to melt and revitrify cryo samples. Optical microscopes are already being used in many cryo-EM groups for correlative light and electron microscopy(de Boer *et al*., 2015). They are also simpler to operate, more cost-effective, as well as more straightforward to adapt to time-resolved experiments, all of which should facilitate their adoption.

## Results

Figure 1A shows a photograph of the optical microscope (Leica DM6000 CFS) that we have adapted for melting and revitrification experiments (Methods). Bright field images are recorded with a CMOS camera placed on top of the instrument, with the sample illuminated with white light from below. The melting laser beam (532 nm, indicated in green) enters the microscope head from the left and is reflected by a dichroic mirror that overlaps it with the optical axis and directs it at the sample. Microsecond laser pulses are obtained by chopping the output of a continuous laser with an acousto-optic modulator. The sample is held by a cryo stage for correlative microscopy (Linkam CMS 196), which maintains it at liquid nitrogen temperature and prevents the condensation of water vapor (Fig. 1B).

**Figure 1.**
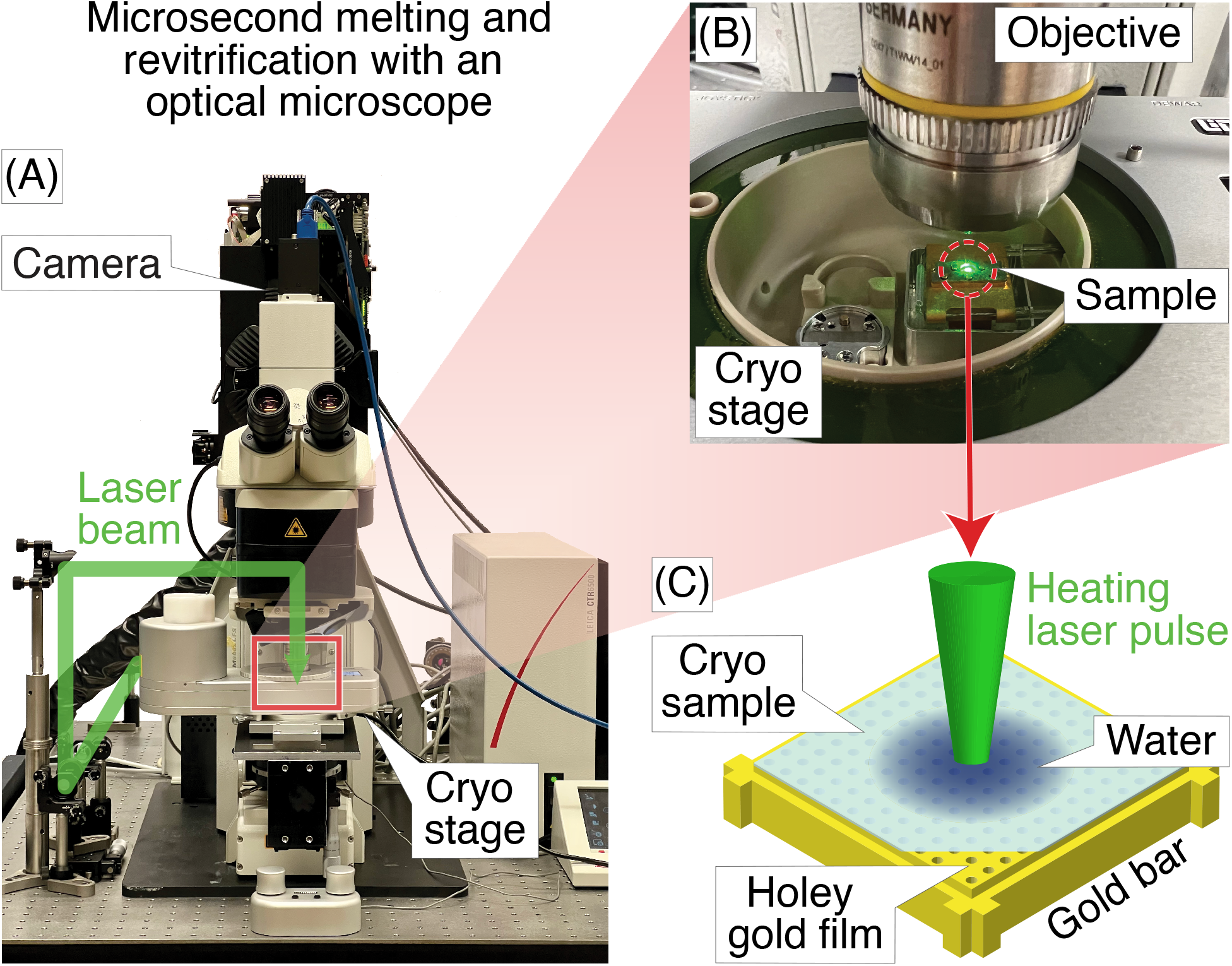
Melting and revitrification of cryo samples with an optical microscope. **(A)** Photograph of the optical microscope and cryo stage. The laser beam (532 nm, indicated in green) enters the head of the microscope from the left and is focused onto the sample. **(B)** Detail of the cryo stage, with the laser beam striking the sample. **(C)** Illustration of the sample geometry. A microsecond laser pulse is used to melt and revitrify the cryo sample in the center of a grid square.

The geometry of a melting and revitrification experiment is illustrated in Figure 1C. The cryo sample is prepared on an UltrAuFoil specimen support, which features a holey gold film on a 300 mesh gold grid. The heating laser is focused onto the center of a grid square (25 μm FWHM beam diameter in the sample plane). Under laser irradiation, the gold film rapidly heats up in the vicinity of the laser beam, causing the cryo sample to melt, so that the embedded particles can undergo unhindered motions in liquid phase. Within a few microseconds, the sample temperature stabilizes near room temperature (Voss *et al*., 2021a). When the laser is switched off, the sample cools within just a few microseconds and revitrifies, trapping the particles in their transient configurations, in which they can subsequently be imaged (Voss *et al*., 2021b). The short cooling time is a consequence of the fact that the laser heats the sample only locally, while its surroundings remain at cryogenic temperature, so that the heat is efficiently dissipated after the end of the laser pulse.

Figure 2 demonstrates that melting and revitrification experiments can be successfully performed with our optical microscope. Figure 2A shows an optical micrograph of a typical grid square of an apoferritin cryo sample. Melting and revitrification with a 40 μs laser pulse (45 mW) barely causes the image contrast to change. However, a characteristic signature of successful melting and revitrification is readily apparent in a difference image (Figure 2B), which is obtained by subtracting a micrograph recorded before revitrification from one recorded after. The revitrified area in the center of the grid square is surrounded by a thin, dark ring and features a dark patch in its center.

**Figure 2.**
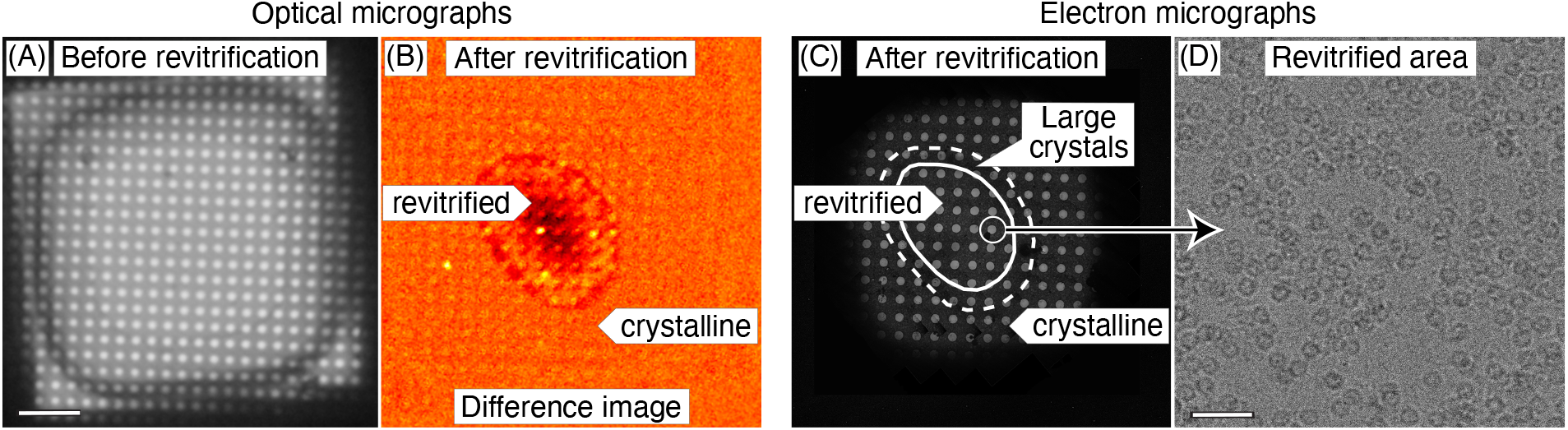
Correlative light-electron microscopy of a cryo sample that was revitrified in the optical microscope. **(A)** Optical micrograph of a typical grid square of an apoferritin cryo sample. **(B)** Difference image of the sample after melting and revitrification with a 40 μs, obtained by subtracting the image before revitrification from one recorded after. The revitrified area in the center of the grid square features a thin, dark outline that arises from the formation of crystals in the surrounding area. A dark patch in the center indicates thinning of the sample due to evaporation. **(C)** An electron micrograph of the same area confirms the interpretation of the image contrast in **(B)**. The outline of the revitrified area is marked with a solid line. A dashed line indicates the region in which the formation of large crystals is observed. **(D)** Micrograph recorded in the area circled in **(C)**, showing intact apoferritin particles. Scale bars, 10 μm in **(A)** and 40 nm in **(D)**.

An electron micrograph of the same area (Figure 2C) confirms the success of the revitrification experiment and elucidates the nature of the features visible in the difference image. The micrograph shows that the sample has melted and revitrified in the vicinity of the laser focus, while it has crystallized in the surrounding areas (details in Supplementary Figure S2). The outline of the revitrified area (solid line) closely agrees with the inner boundary of the thin, dark ring that is visible in the difference image of Figure 2B. This ring seems to originate from the large ice crystals that have formed at the border of the revitrified area (dashed line, see also Supplementary Figure S2 for a magnified view), as previously observed (Voss *et al*., 2021a). The dark spot visible in the center of the revitrified area does not seem to correspond to any feature in the electron micrograph. We propose that it results from the thinning of the sample due to evaporation that occurs during laser heating and which is most pronounced in the center of the laser focus, where the sample reaches the highest temperature (Voss *et al*., 2021a). This interpretation is supported by the fact that the dark spot becomes most pronounced when the sample is evaporated entirely in the center of the grid square (Supplementary Figure S1D).

We conclude that difference images can be used to assess the success of a melting and revitrification experiment on the fly, which can be inferred from the presence of a thin, dark ring that surrounds a dark spot, which marks the center of the laser focus. As illustrated in Supplementary Figure S1, these features are absent if the laser power is too low to induce melting and revitrification. It is also to straightforward to detect if the laser power is too high, so that the sample is evaporated entirely in the vicinity of the laser focus. In this case, a pronounced dark spot results in the center of the grid square, with the holes of the gold film appearing bright (Supplementary Figure S1C, D). In order to ensure that the sample undergoes a reproducible temperature evolution, the diameter of the revitrified area should be kept constant (Voss *et al*., 2021a). This can be achieved by monitoring the diameter of the dark ring in the difference images and adjusting the laser power accordingly.

Melting and revitrification with our optical microscope leaves the proteins intact, as evidenced by an electron micrograph collected from a hole within the revitrified area (Figure 2D). Single-particle reconstructions corroborate this result and show that revitrification does not alter the structure of the apoferritin particles. We revitrified 17 areas of a fresh cryo sample in the optical microscope and imaged them on a Titan Krios G4. A comparable number of images were collected of areas that had not been exposed to the laser beam. The reconstructions from the conventional and revitrified areas feature a resolution of 1.47 Å and 1.63 Å, respectively (Figure 3A). At this resolution, side chain densities are clearly resolved, and individual water molecules can be distinguished (Figure 3B). Within the near-atomic resolution of these reconstructions, the structure of the particles in the conventional and revitrified areas is indistinguishable. We conclude that revitrification in our optical microscope does not alter the protein structure.

**Figure 3.**
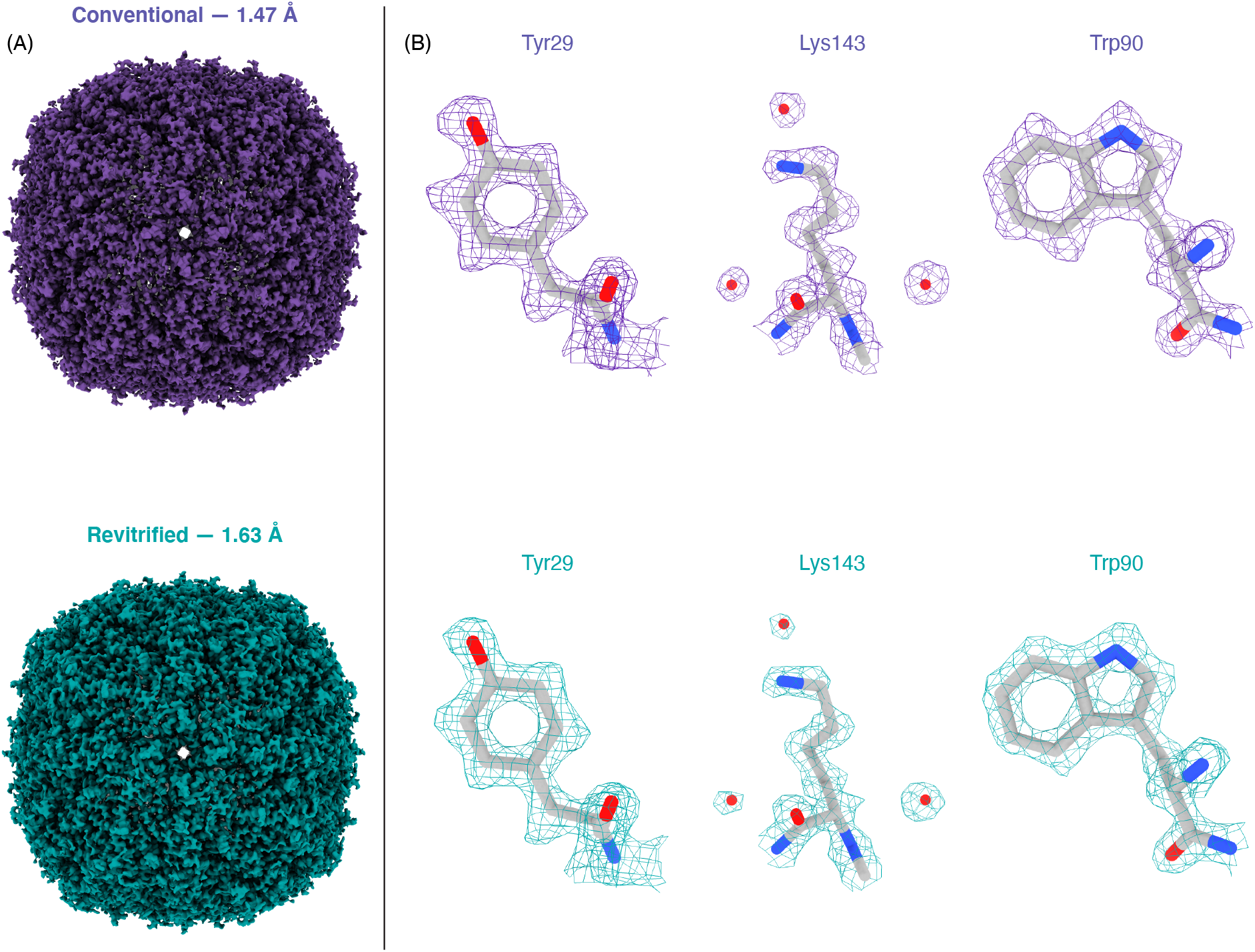
Reconstructions of apoferritin from conventional and revitrified sample areas. Within the spatial resolution of the reconstruction, the structure of the particles is indistinguishable. **(A)** Reconstructions from conventional (purple, 1.47 Å resolution) and from revitrified areas (green, 1.62 Å). **(B)** Details of the reconstructions, showing that the side chain denisities are clearly resolved and that single water molecules are visible. A molecular model of apoferritin (PDB 6v21 (*Wu* et al.*, 2020)*) has been placed into the density through rigid body fitting.

## Conclusions

In summary, we have demonstrated that microsecond time-resolved cryo-EM experiments can be performed with a correlative light-electron microscopy approach. The success of a melting and revitrification experiment can be assessed on the fly by monitoring characteristic signatures in an optical difference image. We have also shown that revitrification in our optical microscope setup preserves the protein structure, allowing us to obtain near-atomic resolution reconstructions. This suggests that the revitrification process does not fundamentally limit the obtainable spatial resolution.

Revitrification with an optical microscope is in several respects complementary to the *in situ* approach we have previously described (Voss *et al*., 2021b, 2021a; Harder *et al*., 2022), with each approach offering distinct advantages. It is usually more straightforward to determine the diameter of the revitrified area in an *in situ* experiment, since the surrounding crystalline region offers a strong contrast in an electron microscope, whereas it may be more difficult to detect in optical difference images if the crystallites are too small. It should however be possible to obtained a better contrast by using different optical imaging modes.

*In situ* experiments also offer the advantage that the behavior of the particles during revitrification can be assessed immediately, either by visual inspection or from an on-the-fly reconstruction, so that the laser parameters can be adjusted as needed. Finally, *in situ* experiments will likely be indispensable to further characterize and advance the experimental approach. For example, we have previously been able to infer that cryo samples partially crystallize during laser melting (Voss *et al*., 2021a). To better understand the phase behavior of cryo samples during rapid heating, we have therefore embarked on time-resolved *in situ* experiments that employ intense high-brightness electron pulses of microsecond duration (Olshin *et al*., 2021) to capture the structural evolution of the cryo sample.

The correlative revitrification approach described here offers the significant advantage that its implementation is technically less involved, which we hope will encourage other research groups to use our technique and explore its potential for elucidating the fast dynamics of a variety of systems. Once suitable experimental conditions for a time-resolved experiment have been established, samples for high-resolution imaging can be most conveniently revitrified with the optical microscope. Because of the simplicity of the setup, the experimental workflow involved can also be easily automated, which we have begun to do. Imaging the sample with the optical microscope also does not damage the particles, in contrast to *in situ* experiments with the electron microscope, where the sample has to be exposed to a small electron dose in order to locate and center areas for revitrification at low magnification. This is significant because the exposure to a dose of only a few electrons/Å^2^ induces so much fragmentation that the proteins will completely unravel once the sample is melted (Voss *et al*., 2021b, 2021a). It is therefore sensible to avoid exposure even to the small dose required for low-magnification imaging, which may potentially cause some proteins to lose their function even though no structural damage is evident (Glaeser, 2016).

Using an optical microscope for revitrification may also enable new types of experiments. The simple optical layout of the instrument makes it straightforward to combine different laser beams and wavelengths, which may be used to trigger different dynamics or monitor the revitrification process. For example, it would be straightforward to use a separate UV laser beam to release a caged compound (Shigeri *et al*., 2001; Ellis-Davies, 2007) in selected grid squares. Once the sample is laser melted, the compound will then become available to initiate protein dynamics (Voss *et al*., 2021b). By releasing different quantities of the caged compound in different grid squares, it would then be possible to study how the concentration of the compound affects the dynamics, with all experiments conducted on the very same specimen grid, thus guaranteeing the most reproducible conditions. It will also become possible to combine revitrification experiments with other types of correlative microscopy. It is even conceivable to perform experiments on entire cells, as long the heat transfer in the sample can be engineered to be fast enough, so that vitrification can be achieved after the end of the laser pulse. Finally, our optical microscope setup may also provide a practical approach to improve the quality of conventional cryo samples. Melting and revitrification can be used to turn some crystalline areas vitreous or reduce the ice thickness through evaporation. It may also be possible to reduce beam induced specimen motion by releasing stress in the vitreous ice film through irradiation with a sequence of laser pulses (Harder *et al*., 2022).

## Methods

Cryo samples were prepared on UltrAuFoil R1.2/1.3 300 mesh grids (Quantifoil), which were rendered hydrophilic through plasma cleaning for 1 min (Tedpella “Easy glow” discharge system, negative polarity with a plasma current of 0.8 mA and a residual air pressure of 0.2 mbar). A volume of 3 μl of the sample solution (mouse heavy chain apoferritin, 8.5 mg/ml in 20 mM HEPES buffer with 300 mM sodium chloride at pH 7.5) was applied to the specimen grid, which was then plunge-frozen with a Vitrobot Mark IV (Thermo Fisher Scientific, 3 s blotting time, 95% relative humidity at a temperature of 10 °C). The grids were screened on a JEOL 2200FS (Olshin *et al*., 2020) before revitrifying them in the optical microscope.

Revitrification experiments were performed with a modified Leica DM6000 CFS optical microscope equipped with a Linkam CMS 196 cryo stage (Figure 1A). Microsecond laser pulses for melting and revitrification are obtained by chopping the output of a 532 nm continuous wave laser (Laser Quantum, Ventus 532) with an acousto-optic modulator (AA-optoelectronic). As shown in Figure 1A, the laser beam enters the microscope head from the left and is reflected by a dichroic mirror that overlaps it with the optical axis. The beam is focused to a spot size of 25 μm FWHM in the sample plane, as determined from an image of the beam recorded with a CCD camera placed in the sample location. Optical bright field images are recorded with a Teledyne FLIR Grasshopper3 camera. Difference images for assessing the success of a revitrification experiment (Figure 2B) were obtained by subtracting an image of the sample before laser irradiation from one recorded after. Both images were acquired as averages of 15 exposures each (1 s), which were aligned using phase correlation-based image registration (Reddy and Chatterji, 1996).

The electron micrographs of Figure 2C, D were recorded on a JEOL 2200FS (Olshin *et al*., 2020). The image in Figure 2C was obtained by stitching together several micrographs acquired at higher magnification, in which the crystalline areas can be more readily identified. Micrographs for single-particle reconstructions (Figure 3) were acquired with a Titan Krios G4 (Thermo Fisher Scientific) operated at 300 kV accelerating voltage and using a 10 eV slit width (Selectris X energy filter). The micrographs were recorded with a Falcon 4 camera, with an exposure time of about 2.5 s and a total dose of 50 electrons Å^-2^. The pixel size was 0.455 Å, and defocus values were in the range of 0.9–1 μm.

Single-particle reconstructions were performed in CryoSPARC 3.3.2 (Punjani *et al*., 2017), as detailed in Supplementary Material 1. Briefly, the conventional (revitrified) apoferritin dataset comprises 10145 (12047) images. Patch motion correction and CTF estimation yielded 6242 (3671) images with a resolution better than 6 Å, which were kept for further processing. Using template-based particle picking, 2937383 (1714924) particles were identified. Following two rounds of 2D classification, 534606 (212377) particles were retained for *ab initio* reconstruction (*C1* symmetry), followed by heterogeneous refinement (*O* symmetry) using three classes. The 457940 (183700) particles found in the most populated classes were then used for homogeneous refinement (*O* symmetry) to give a final map with a resolution of 1.47 Å (1.63 Å).

Reconstructions were visualized with Chimera X (Goddard *et al*., 2018), using contour levels of 0.077 and 0.24 in Figure 3A and Figure 3B, respectively. The molecular model displayed in Figure 3B (PDB 6v21 (Wu *et al*., 2020)) was placed into the map using rigid body fitting.

## Associated Content

### Supplementary Material

Details of the workflow for single-particle reconstructions

Supplementary Figures 1–3

### Data Availability Statement

The data that support the findings of this study are available from the corresponding author upon reasonable request. The maps of apoferritin obtained from conventional and revitrified sample areas have been deposited on EMDB (EMD-15619 and EMD-15620, respectively), and the corresponding data sets on EMPIAR (EMPIAR-11212 and EMPIAR-11213).

### Author Information

**Corresponding author:** Ulrich J. Lorenz

**Notes:** The authors declare no competing financial interests.

## Acknowledgements

This work was supported by the ERC Starting Grant 759145 and by the Swiss National Science Foundation Grant PP00P2_163681. We acknowledge the support of the Dubochet Center for Imaging (DCI) in Lausanne.

