## Supplementary Information for "Microsecond melting and revitrification of cryo samples with a correlative light-electron microscopy approach"

**Supplementary Material 1. Single-particle reconstructions**

The conventional (revitrified) apoferritin dataset consists of 10145 (12047) movies in EER format, which were processed with CryoSPARC 3.3.2 (Punjani *et al.*, 2017), using 40 fractions and no upsampling. Patch motion correction and CTF estimation were performed on all videos with default settings, yielding 6242 (3671) micrographs with a resolution below 6 Å and a relative ice thickness between 1.0 and 1.1, which were kept for further processing.

The following procedure was used to perform template-based particle picking on both data sets. First, blob picking was applied to the remaining images of the conventional dataset, using a radius between 110 and 120 Å. The 881837 particles found were then filtered based on the NCC and power score. The remaining 596829 particles were extracted with a box size of 560 pixels and Fourier cropped to 256 pixels. After 2D classification with 50 classes, the 12 classes of highest quality were manually selected (222088 particles). Using these classes for template-based particle picking yielded 2937383 (1714924) particles for the conventional (revitrified) dataset. Particles with an NCC score above 0.34 were then extracted with a box size of 560 pixels and Fourier cropped to 256 pixels, yielding a total of 727135 (355918) particles.

Reconstructions were obtained with the following procedure. The selected particles were subjected to 2D classification with 50 classes, and the best 29 (18) classes were manually selected. The remaining 645679 (264590) particles were subjected to a second round of 2D classification with 30 classes (150 Å circular mask), of which the best 11 (9) classes were manually selected. The remaining 534606 (212377) particles were used for *ab-initio* reconstruction with *C1* symmetry and 3 classes. Subsequently, two rounds of heterogeneous refinement were performed with 3 classes, using *C1* symmetry in the first iteration and *O* symmetry in the second. After each iteration, the scarcely populated classes were rejected. The remaining particles were then re-extracted with a box size of 600 pixels (without Fourier cropping), leaving 451420 (957860) particles. Homogeneous refinement with *O* symmetry, using per-particle defocus optimization, per-group CTF refinement, correction of magnification anisotropies, and Ewald sphere correction yielded a 1.47 Å (1.63 Å) resolution map (Gold Standard FSC at 0.143).

### Supplementary Figures

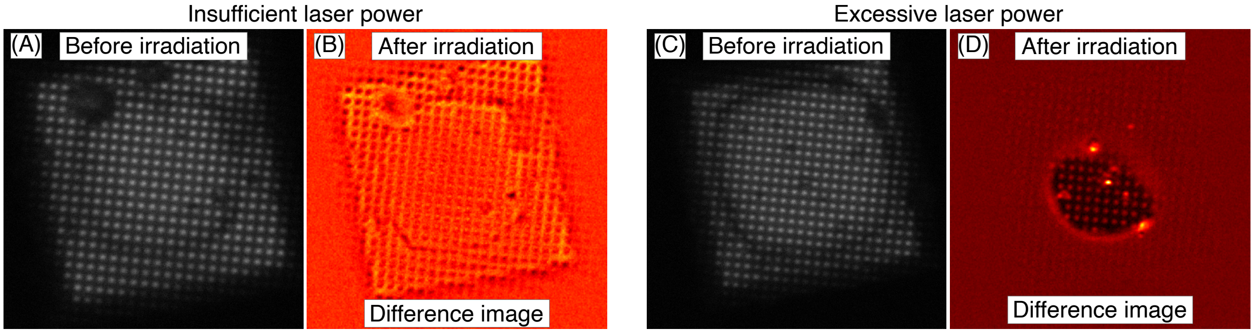


**Supplementary Figure S1. Optical micrographs of unsuccessful revitrification experiments. (A, B)** Insufficient laser power. The difference image **(B)** does not exhibit the characteristic contrast changes associated with successful melting and revitrification that can be seen in Fig. 2**B**. Most features in the difference image correspond to structures visible before laser irradiation **(A)** and therefore likely arise from a change in defocus due to a deformation of the specimen grid under laser irradiation. **(C, D)** Excessive laser power. The sample evaporates in the center of the grid square, which leads to strong contrast changes in the difference image, with the holey gold film appearing dark in the evaporated area, and the holes bright.

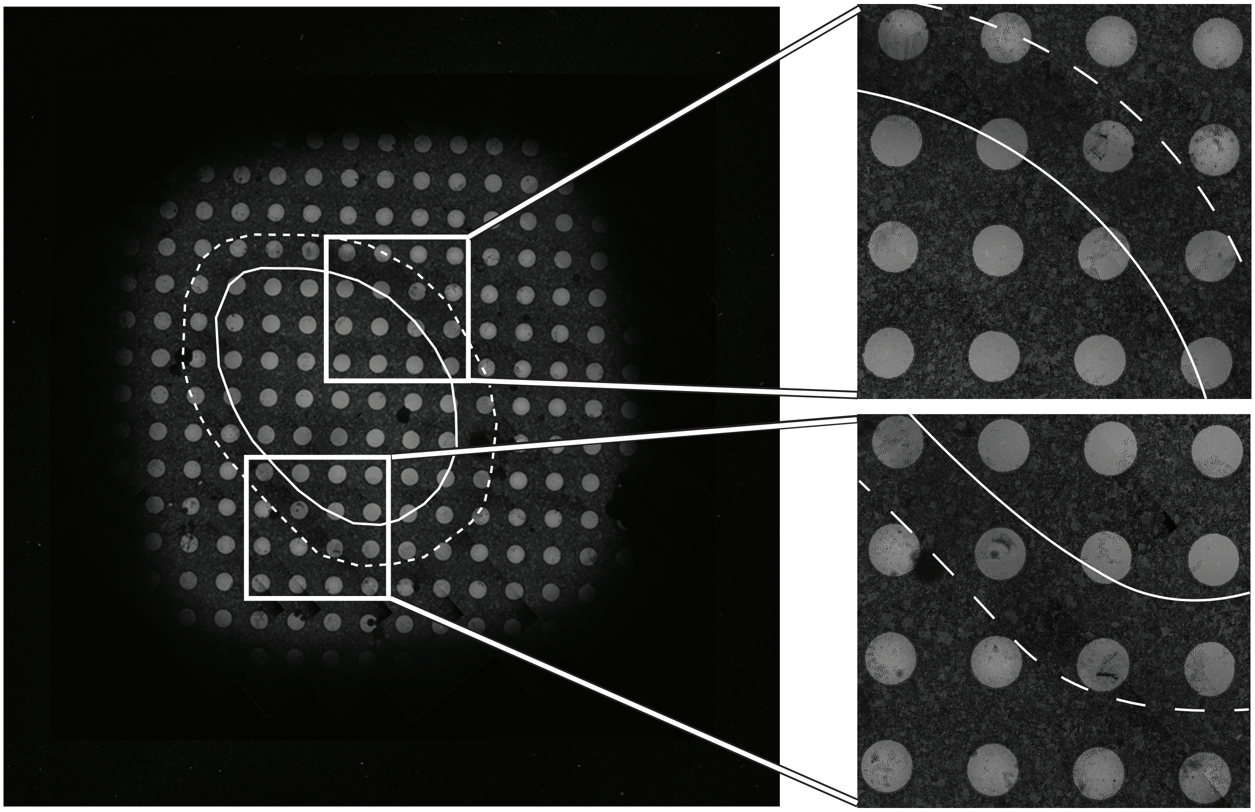

**Supplementary Figure S1. Electron micrograph of the revitrified cryo sample from Fig. 2(C) with details of the boundary between the revitrified and crystalline areas**. The outline of the revitrified area is marked with a solid line. A dashed line indicates the region in which the formation of large crystals is observed, which causes a thin, dark outline to appear in the optical difference image of Fig. 2**(B)**.

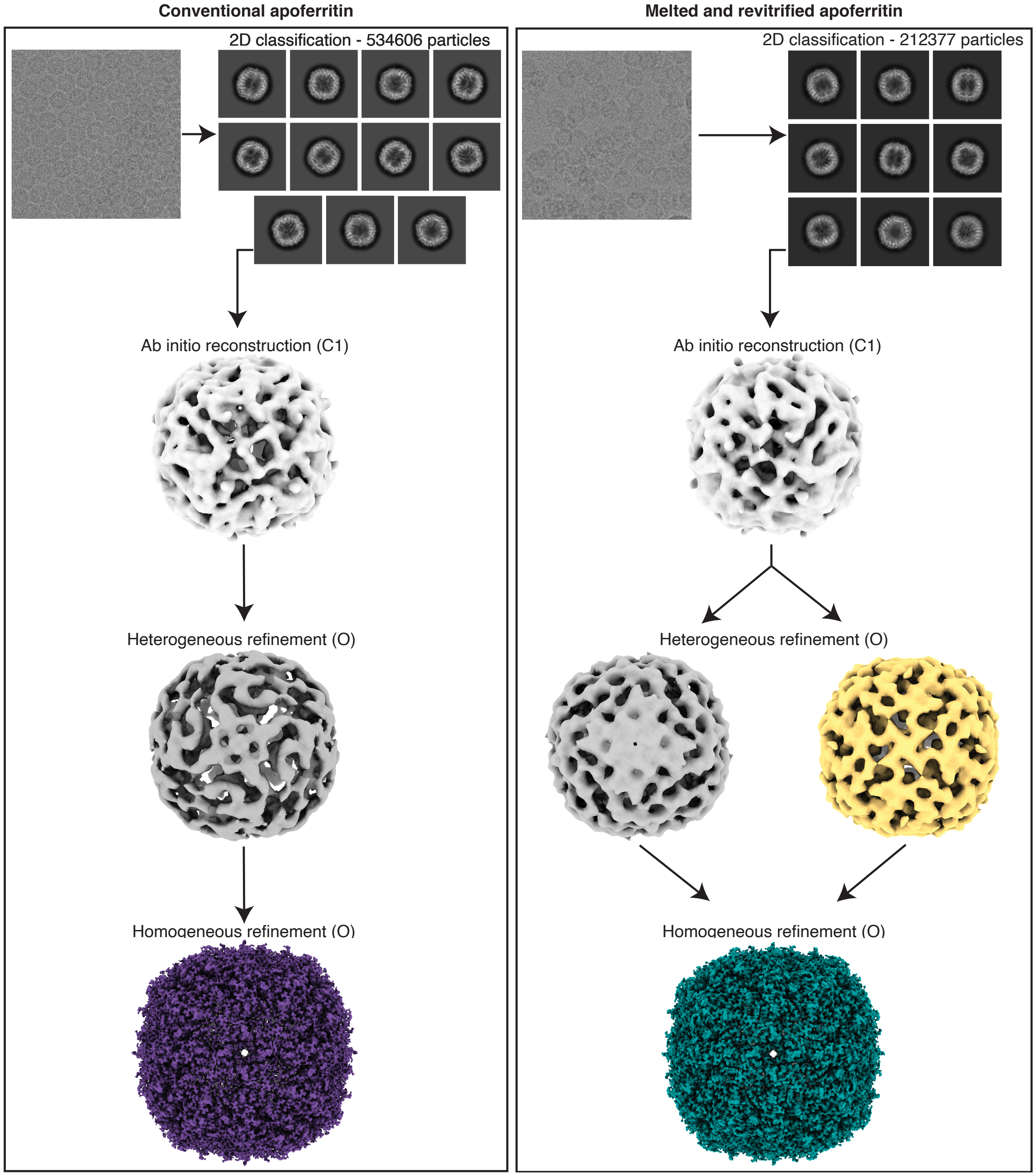


**Supplementary Figure S3. Workflow for the single-particle reconstructions of apoferritin from conventional and revitrified sample areas.**
